# Editing of the urease gene by CRISPR-Cas in the diatom *Thalassiosira pseudonana*

**DOI:** 10.1101/062026

**Authors:** Amanda Hopes, Vladimir Nekrasov, Sophien Kamoun, Thomas Mock

## Abstract

**Background**: CRISPR-Cas is a recent and powerful edition to the molecular toolbox which allows programmable genome editing. It has been used to modify genes in a wide variety of organisms, but only two alga to date. Here we present a methodology to edit the genome of *T. pseudonana*, a model centric diatom with both ecological significance and high biotechnological potential, using CRISPR-Cas.

**Results**: A single construct wa assembled using Golden Gate cloning. Two sgRNAs were used to introduce a precise 37nt deletion early in the coding region of the urease gene. A high percentage of bi-allelic mutations (≤ 61.5%) were observed in clones with the CRISPR-Cas construct. Growth of bi-allelic mutants in urea led to a significant reduction in growth rate and cell size compared to growth in nitrate.

**Conclusions**: CRISPR-Cas can precisely and efficiently edit the genome of *T. pseudonana*. The use of Golden Gate cloning to assemble CRISPR-Cas constructs gives additional flexibility to the CRISPR-Cas method and facilitates modifications to target alternative genes or species.

## Background

Diatoms are ecologically important microalgae with high biotechnological potential. Since their appearance about 240 million years ago [1], they have spread and diversified to occupy a wide range of niches across both marine and freshwater habitats. Diatom genomes have been shaped by secondary endosymbiosis and horizontal gene transfer resulting in genes derived from heterotrophic hosts, autotrophic endosymbionts and bacteria [2, 3]. They play a key role in carbon cycling [4], the food chain, oil deposition and account for about 20% of the world’s annual primary production [5, 6]. However, they are perhaps best known for their intricate silica frustules which give diatoms a range of ecological advantages and play a key role for carbon sequestration and silica deposition.

Several aspects of diatom physiology including the silica frustule, lipid storage and photosynthesis are being applied to biotechnology. Areas of high interest include nanotechnology [7], drug delivery [8], biofuels [9], solar capture [10] and bioactive compounds [11].

Given the ecological importance of diatoms and their applications for biotechnology, it is pivotal that the necessary tools are available to study and manipulate them at a molecular level. This includes the ability to replace, tag, edit and impair genes. A recent edition to the genetic tool box, CRISPR-Cas, allows double strand breaks (DSBs) to be introduced at specific target sequences. This adapted mechanism, used by bacteria and archaea in nature as a defence system against viruses, facilitates knock-out by the introduction of mutations through repair by error prone non-homologous end joining (NHEJ) or homologous recombination (HR). This requires both a Cas9 to cut the DNA and a sgRNA to guide it to a specific sequence. Further information on the history and application of CRISPR-Cas can be found in several excellent reviews [12–14]. Zinc-finger nucleases (ZFNs), meganucleases and transcription activator-like effector nucleases (TALENs) have also been used to induce double strand breaks. TALENs and CRISPR-Cas both bring flexibility and specificity to gene editing, however CRISPR-Cas is also cheap, efficient and easily adapted to different sequences by simply changing the 20nt guide sequence in the sgRNA.

So far, within the diverse group of algae, the diploid, pennate diatom *Phaeodactylum tricornutum* [15] and the haploid, green alga *Chlamydomonas reinhardtii* [16] have been subject to gene editing by CRISPR-Cas. NHEJ and HR have been used to repair DSBs following CRISPR-Cas or TALENs in *P. tricornutum*, introducing mutations into a nuclear coded chloroplast signal recognition particle [15], the urease gene [17] and several genes associated with lipid metabolism [18]. *Thalassiosira pseudonana* is a logical choice for CRISPR-Cas development. It is a model centric diatom with a sequenced genome (first eukaryotic marine phytoplankton to be sequenced [2]) and well-established transformation systems [19, 20]. The genus has multiple biotechnology applications [8, 21, 22], and although gene silencing has been established, a method to easily and efficiently knockout and edit genes and the entire genome would be highly advantageous. The genus *Thalassiosira* is among the top 10 genera of diatoms in the World’s Ocean in terms of ribotype (V9 of 18S) diversity and abundance [23] and the species *T. pseudonana* is a model for understanding the mechanisms behind silicification [24–26].

Golden Gate cloning can add further flexibility to CRISPR-Cas methods as demonstrated in higher plants [27]. As a modular cloning system it allows different modules, including the sgRNA, to be easily interchanged or added [28]. As a result, new constructs can be made quickly, cheaply and efficiently for new or multiple targets. This extends to any aspect of the construct, including promoters, Cas9 variants and their nuclear localisation signals (NLS). As a result, construct alterations such as replacing constitutive promoters for inducible ones, exchanging the wildtype Cas9 for a Cas9 nickase or changing the localisation signal to target other organelles can be easily carried out.

An increasing range of software tools are available for CRISPR-Cas, including programs that facilitate sgRNA target searches in a genetic locus of interest, estimate efficiencies of sgRNAs [29] and perform off-target predictions.

While off-target prediction tools tend to be species specific, there are tools that accept requests for a genome to be added to the list, or allow for a genome to be directly uploaded [30, 31]. The latter is particularly useful for less studied organisms, such as diatoms. The ability to combine several different aspects of sgRNA design can help to make an informed decision when choosing target sites for gene editing.

Our paper represents a proof of concept to demonstrate the feasibility of gene editing in the model diatom *T. pseudonana* using two sgRNAs to induce a precise deletion in the urease gene. Methods combine a flexible Golden Gate cloning approach with sgRNA design, which draws on several available online tools. This takes into account multiple factors, such as position within the gene in terms of both early protein disruption and presence in the coding region, DNA cutting efficiency and presence of restriction enzyme sites at the cut site. The latter, in combination with inducing a large deletion by targeting with two sgRNAs, allows easy screening of mutants through either the restriction enzyme site loss assay [32] or the PCR band-shift assay [33], respectively.

## Method

### Strains and growth conditions

*Thalassiosira pseudonana* (CCMP 1335) was grown in 24h light (100-140 μE) at 20°C in half salinity Aquil synthetic seawater [34]. For routine growth, a 1mM nitrate concentration was used.

### 5’ RACE U6 promoter

To identify the U6 small nuclear RNA (snRNA) in *T. pseudonana*, an NCBI blastn search was performed on the genome against the central conserved region of the U6 sequence. Two potential guanine (G) start sites were found downstream of a TATA box in the promoter. To identify the start site of the U6 snRNA and empirically determine the end of the promoter, 5’ RACE was carried out as follows: 400ml of culture was grown to exponential phase (1×10^6^ cells ml^−1^) and harvested. Small RNAs were extracted and enriched using a miRNeasy kit (Qiagen). 5’ template switching oligo RACE was performed according to Pinto and Lindblad [35]. For oligos used see Table 1 (ref. numbers 1–3). RACE products were sequenced and results aligned to the genome to determine the end of the promoter.

**Table 1.**
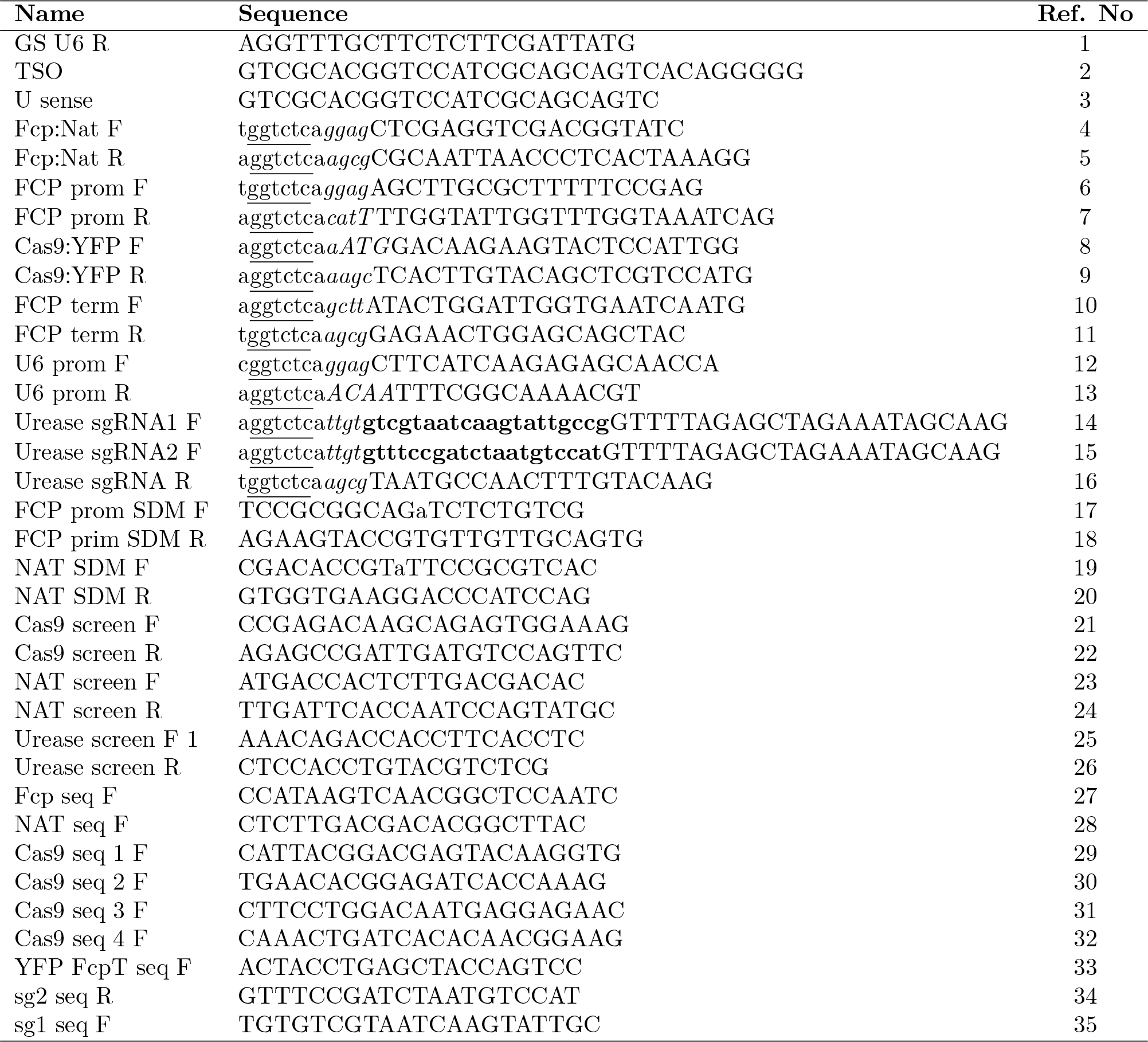
Oligonucleotides used in this study. Ref. N° 1–3: oligos used in 5’ RACE [35]. Ref. N° 4–16: primers for Golden Gate cloning, BsaI sites are <u>underlined</u>, 4nt overhangs are shown in *italics*, and sgRNA targets are shown in **bold.** Upper case indicates complement to the template. Ref. N° 17–20: primers for SDM, lower case indicates base change. Ref. N° 21–26: primers for screening transformants. Ref. N° 27–35: primers for sequencing the CRlSPR-Cas construct.

### Plasmid construction using Golden Gate cloning

Golden Gate cloning was carried out according to Weber et al. [28] and Belhaj et al. [33]. Bsal and Bpil sites were removed in a so-called “domestication” procedure using a Q5 site-directed mutagenesis (SDM) kit (NEB). For oligos used in SDM see Table 1 (ref. numbers 17–20). BsaI sites and specific 4nt overhangs for Level 1 (L1) assembly were added through PCR primers (Table 1). Plasmid DNA was extracted using a Promega mini-prep kit.

#### Golden Gate reactions

Golden Gate reactions for L1 and Level 2 (L2) assembly were carried out using the method specified in Weber et al. [28]. Forty fmol of each component was included in a 20μl reaction with 10 units of Bsal or Bpil and 10 units of T4 DNA ligase in 1× ligation buffer. The reaction was incubated at 37°C for 5 hours, 50°C for 5 minutes and 80°C for 10 minutes. Five μl of the reaction was transformed into 50μl of NEB 5-alpha chemically competent *E. coli*.

#### Level 0 assembly

The endogenous FCP promoter and terminator were amplified with GoTaq flexi (Promega) from domesticated pTpFCP/NAT [19] and the U6 promoter from gDNA (extracted with an Easy-DNA gDNA purification kit (Thermo Fisher)). Both promoters are associated with high expression levels. The U6 promoter was amplified from the position −470 to −1 (the end of the promoter), cutting off a BpiI site and removing the need for additional SDM. For oligos, see Table 1 (ref. numbers 6–7 and 10–13). Products were cloned into a pCR8/GW/TOPO vector (Thermo Fisher).

Domesticated human codon bias Cas9 from *Streptococcus pyogenes* with an N-terminal SV40 NLS and a C-terminal YFP tag was PCR-amplified using Phusion DNA polymerase (NEB) and L1 Cas9:YFP plasmid as a template. The PCR product was purified with a GFX PCR DNA and gel purification kit (GE Healthcare) and incubated for 20 minutes with Taq to add adenine overhangs before cloning directly into a pCR8/GW/TOPO vector. For oligos, see Table 1 (ref. numbers 8–9).

#### Level 1 assembly

The FCP:NAT cassette was PCR-amplified using Phusion polymerase and the domesticated pTpFCP/NAT as a template, purified and inserted into a L1 pICH47732 destination vector. FCP promoter, Cas9 and FCP terminator L0 modules were assembled into L1 pICH47742. For oligos, see Table 1 (ref. numbers 4–5).

The sgRNA scaffold was amplified from pICH86966_AtU6p_sgRNA_NbPDS [32] with sgRNA guide sequences integrated through the forward primers. Together with the L0 U6 promoter, sgRNA_Urease 1 and sgRNA_Urease 2 were assembled into L1 destination vectors pICH47751 and pICH47761, respectively. For oligos, see Table 1 (ref. numbers 14–16).

#### Level 2 assembly

L1 modules pICH47732:FCP:NAT, pICH47742:FCP:Cas9YFP, pICH47751:U6:sgRNA_Urease 1, pICH47761:U6:sgRNA_Urease 2 and the L4E linker pICH41780 were assembled into the L2 destination vector pAGM4723. Constructs were screened by digestion with EcoRV and sequenced. For oligos used in sequencing, see Table 1 (ref. numbers 27–35). See Figure 1 for an overview of the Golden Gate assembly procedure and the final construct.

**Figure 1.**
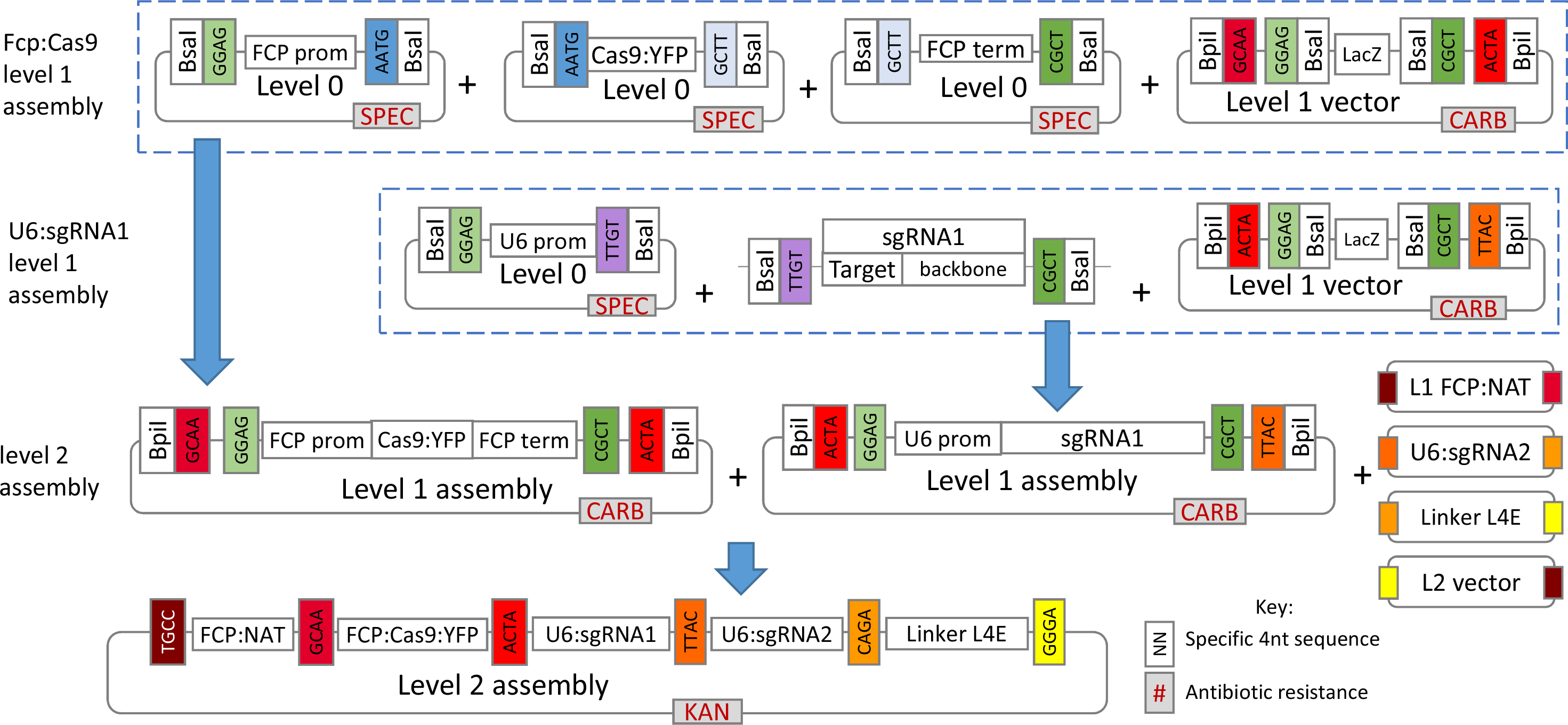
Overview of level 1 and level 2 Golden Gate cloning for assembly of the CRlSPR-Cas construct pAGM4723:TpCC_Urease. Level 1 assemblies of pICH47742:FCP:Cas9YFP and plCH47751:U6:sgRNA_Urease 1 are shown. Bsal or Bpil restriction enzymes cut outside the recognition site leading to specific 4nt overhangs which are complementary to adjacent modules, allowing several modules to be accurately assembled in one reaction. Complementary 4nt sequences are colour coded to indicate adjacent modules.

### sgRNA design for the urease gene knockout

Two sgRNAs were designed to cut 37nt apart early in the coding region of the urease gene (JGI ID 30193) to induce a deletion and frame-shift. Several programmes, explained below, were used to collect data and make an informed decision on sgRNA choice. Excel was used to combine, process and compare data.

#### Selecting CRISPR-Cas targets and estimating on-target score

Twenty bp targets with an NGG PAM were identified and scored for on-target efficiency using the Broad Institute sgRNA design programme (www.broadinstitute.org/rnai/public/analysis-tools/sgrna-design), which utilises the Doench et al. [29] on-target scoring algorithm calculated from >1800 empirically tested sgRNAs.

#### Determining cut positions and cross referencing to restriction recognition sites

All restriction sites and their positions within the urease gene were identified using the Emboss restriction tool (http://emboss.bioinformatics.nl/). As the Broad Institute sgRNA design programme does not give the location of CRISPR-Cas targets within a gene, this was determined using Primer map (http://www.bioinformatics.org/sms2/primer_map.html [36]). The cut site position (3nt upstream of the start of the PAM sequence) was calculated for each sgRNA depending on sense or anti-sense strand placement. All predicted CRISPR-Cas cut sites were cross-referenced to restriction recognition sites.

#### Reverse complement of antisense strand CRlSPR-Cas targets

The reverse complement (RC) was found for each CRlSPR-Cas target using the programme: http://www.bioinformatics.org/sms2/rev_comp.html [36]. In the final spreadsheet (Supplementary Figure 1), if a target was located on the anti-sense strand, the RC was shown for the ‘sense strand sequence’ column. This allows the sgRNA to be easily searched within the original gene sequence.

#### Determine position of CRlSPR-Cas cut sites in relation to coding region

An array was made with start and end positions for each exon/intron. Cut site positions were compared to exon/intron ranges and the relevant exon/intron returned if the data overlapped. The final spreadsheet gives data on CRlSPR-Cas target sequences and their sense sequence (if located on the antisense strand), location of target (relative to the sense strand), predicted CRlSPR-Cas cut site, first nucleotide of the target, PAM sequence, location (i.e. exon, intron), strand, sgRNA score and restriction recognition sites overlapping the cut site. The table (Supplementary Figure 1) was sorted to prioritise sgRNAs by starting base prioritising guanine, sgRNA score, position within the gene and interaction with restriction recognition sites.

#### Predicting off-targets

The full 20nt target sequences and their 3’ 12nt seed sequences were subjected to a nucleotide BLAST search against the *T. pseudonana* genome. Resulting homologous sequences were checked for presence of an adjacent NGG PAM sequence at the 3’ end. The 8nt sequence outside of the seed sequence was manually checked for complementarity to the target sequence. In order for a site to be considered a potential off-target the seed sequence had to match, a PAM had to be present at the 3’ end of the sequence and a maximum of three mismatches between the target and sequences from the blast search were allowed outside of the seed sequence.

Off-targets were also checked using the EuPaGDT program [31], which checks for up to 5 mismatches in the 20nt target sequence and the CasOT program [30], which uses flexible parameters for identifying off-target sequences. Parameters were set to check for an NGG PAM, complete complementarity within the 12nt seed sequence and up to 3 mismatches outside of the seed region.

### Transformation and selection

Using the Poulsen et al. [19] method, transformations were carried out in triplicate with the CRlSPR-Cas construct, pTpfcp/nat (positive control) and water (negative control). Five × 10^7^ cells in exponential phase were used per shot with a rupture disc of 1350psi and a 7cm flight distance. Following transformation, cells were rinsed into 25ml of media and left to recover for 24 hours under standard growth conditions. Cells were counted using a Coulter counter (Beckman) and 2.5 × 10^7^ cells from each transformation were spread onto 5, ½ salinity Aquil 0.8% agar plates (5 × 10^6^ cells/ plate) with 100μg ml^−1^ nourseothricin. Plates were incubated under standard conditions for two weeks. Remaining sample was diluted to 1 × 10^6^ cell ml^−1^ in media and supplemented with nourseothricin to final a concentration of 100μg ml^−1^ for liquid selection. Liquid selection cultures were maintained under standard growth conditions with 100μg ml^−1^ nourseothricin. Colonies were picked and transferred to 20μl of media. Ten μl from each colony was transferred to 1ml of selective media for further growth. The remaining sample was used in screening.

To isolate sub-clones from colonies which screened positive for mutations, 100μl of cells at exponential phase were streaked onto ½ salinity Aquil 0.8% agar plates with 100μg ml^−1^ nourseothricin.

### Screening clones and cultures

Ten μl from each colony or culture from liquid selection, was spun down and supernatant removed. Cells were re-suspended in 20μl of lysis buffer (10% Triton X-100, 20mM Tris-HCl pH8, 10mM EDTA), kept on ice for 15 minutes then incubated at 95°C for 10 minutes. One μl of lysate was used in Taq PCR to amplify the CRISPR-Cas targeted fragment of the urease gene. Clones were also screened for Cas9 and NAT. For PCR primers, see Table 1 (ref. numbers 21– 26). PCR products were run on an agarose gel to check for the lower MW band associated with a double-cut deletion in the urease gene and for the presence of Cas9 and NAT. Urease PCR products were also digested with BsaI and HpaII to determine if the restriction recognition sites, which overlap the cut sites, had been mutated. PCR products were sent for sequencing to confirm mutations.

### Growth experiments

Knockout and wild-type (WT) cultures were nitrate depleted by growing cells in nitrate free media until cell division stopped and quantum yield of photosynthesis (Fv/Fm measured on the Phyto-PAM-ED) dropped below 0.2. Cultures were then transferred in triplicate at a final concentration of 2.5 × 10^4^ cells ml ^−1^ into 25ml of media with either 1mM sodium nitrate or 0.5mM urea. Cell count and mean cell size were measured once a day using a Coulter counter. Fv/Fm measurements were also taken daily. Growth rates were calculated using μ= Ln_2_−Ln_1_/ T_2_−T_1_, where T is a time point corresponding to exponential growth and Ln is the natural log of cell counts ml^−1^. Analysis of variance with Tukey’s pairwise comparision was used to compare both growth rates and cell size at the end of exponential phase between samples.

## Results and discussion

### sgRNA design

The two CRISPR-Cas targets with the highest on target scores (0.5 and 0.79), containing a predicted cut site over a restriction site and occurring early in the coding region, were chosen. sgRNAs were designed to cut 37nt apart at positions 138 and 175 within the urease gene. Both targets started with a G for polymerase III transcription (Figure 2). No off-target sites were predicted for sgRNAs designed for either of the two CRISPR-Cas target sequences.

**Figure 2.**
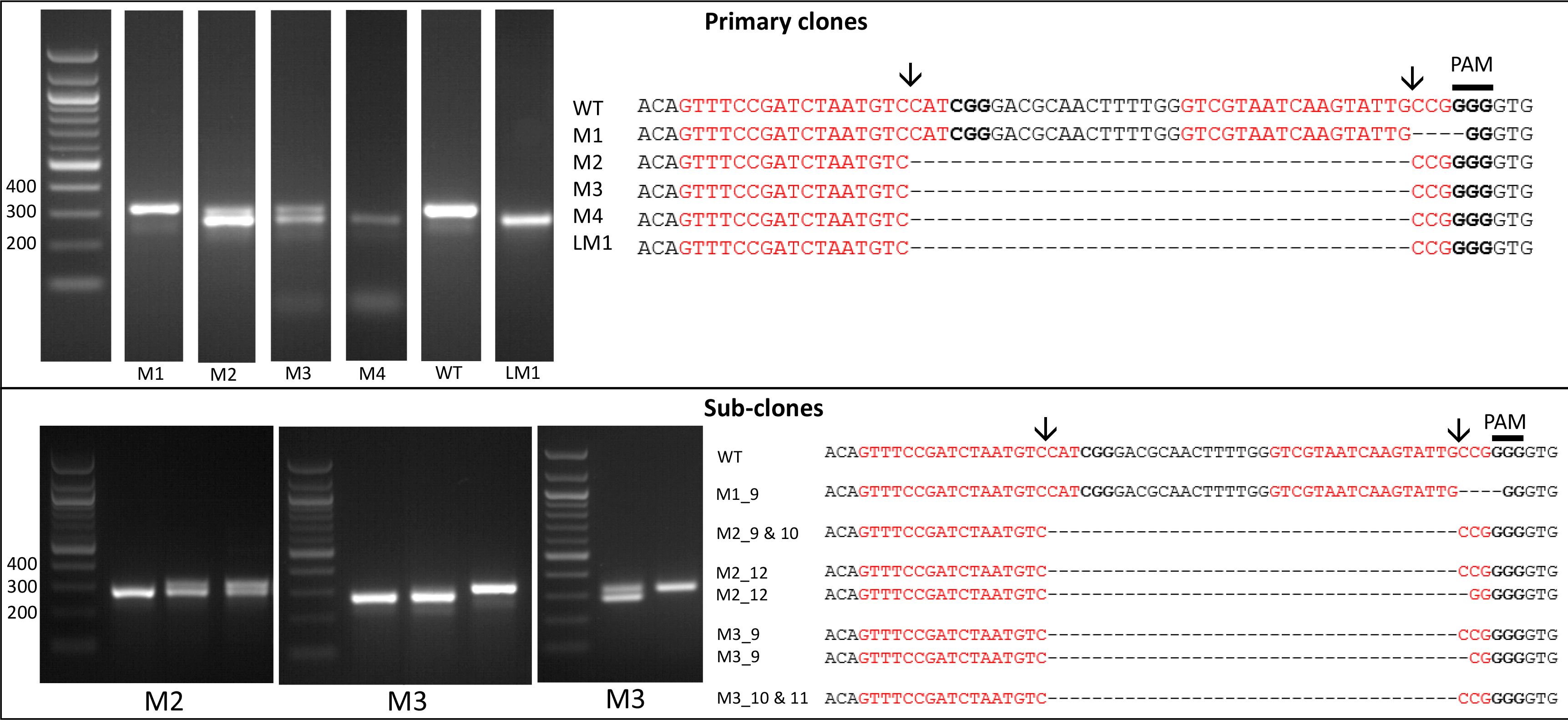
Screening by PCR and sequencing. Expected sgRNA cut indicated by ↓. PCR of targeted urease fragments from primary clones show a single higher MW band for M1 and WT, a lower MW band associated with the expected 37nt deletion for M4 and LM1 and two bands for M2 & M3. Sequence alignments of urease products with mutations are shown for primary clones. A few examples of PCR products from sub-clones are shown. Primary clones M2 and M3 appear to be mosaic with sub-clones containing bi-allelic and mixed products corresponding to full length urease and the lower MW band associated with the deletion. Sequence alignments from mono-allelic subclone M1_9 and bi-allelic sub-clones from M2 and M3 are shown.

### Constructing the CRISPR-Cas plasmid using the Golden Gate cloning method

A single CRISPR-Cas construct was made using Golden Gate cloning (Figure 1). The construct included the NAT selectable marker and Cas9:YFP driven by an endogenous FCP promoter for high expression and two U6 promoter-driven sgRNAs. RNA polymerase III U6 promoters are a popular choice for expression of sgRNAs in CRISPR-Cas [15, 27, 37–39]. RACE products showed that the U6 promoter ended 23nt after the TATA box. As a standardised, efficient, modular system, Golden Gate cloning gives a high level of flexibility to the CRISPR-Cas method and bypasses the need for cotransformation as it enables assembly of multiple expression units, such as Cas9 and sgRNAs, into a single vector backbone. Multiple sgRNA modules can be incorporated into the construct to target several genes or whole pathways. In human cells, up to 7 sgRNAs have been successfully assembled and expressed from a single construct created using the Golden Gate cloning method [39]. Golden Gate has also proved successful for building constructs for genome editing in higher plants using both TALENs [41] and CRISPR-Cas [33, 37].

In this study, only the promoters and target sequences are specific to *T. pseudonana*, which demonstrates how simple it can be to apply this method to a new species using the Golden Gate system. The *S. pyrogenes* Cas9 with a human codon bias, shown previously to work in higher plants [32, 33, 37], carries a SV40 NLS, which follows a canonical sequence found throughout eukaryotes, including *T. pseudonana*.

The long term effects from off-target mutations introduced through CRISPR-Cas are currently unknown, therefore it may be advantageous for future work to remove CRISPR-Cas constructs from mutants. Adding a yeast CEN6-ARSH4-HIS3 sequence to plasmids allows autonomous replication in diatoms and expression of genes without random integration into the genome [20]. Furthermore, removing selection leads to plasmids being discarded. By expressing CRISPR-Cas genes and selective markers on a removable episome, mutations could be introduced without integration of the plasmid. CRISPR-Cas constructs could then be expelled by removing selection. As well as considerations for long term off-target effects, this could also be advantageous for studies and applications which are sensitive to the presence of transgenes.

### Selecting and screening for mutations in the urease gene

The transformation efficiency with the CRISPR-Cas construct was on average 41.5 colonies μg^−1^ plasmid (13.35-66.65 colonies μg^−1^ plasmid). Thirty three colonies were screened by PCR and sequencing of the targeted urease gene fragment.

Four colonies showed mutations in the urease gene. All colonies screened positive for NAT but only the four colonies with mutations screened positive for Cas9, suggesting that once the Cas9 and sgRNAs are present there is a high chance of inducing mutations in the target gene. The lack of Cas9, which accounts for a third of the construct, in the majority of colonies was potentially caused by shearing of the plasmid during microparticle bombardment [38] from either mechanical force or chemical breakdown [42].

Of the four primary colonies which screened positive for mutations (Figure 2), one (M4) showed a single band with a 37nt deletion between the two sgRNA cut sites which suggests that both copies of the urease gene contain the deletion giving a bi-allelic mutant. Two colonies (M2 and M3) produced two bands following PCR: a WT higher MW band and a lower MW band with the 37nt deletion, confirmed by sequencing (Figure 2). The fourth colony (M1) showed a single band associated with the WT urease, however sequencing showed two products: a WT urease and a mutant urease with a 4nt deletion at the first sgRNA cut site. A mixture of PCR products may be due to a mono-allelic mutation, in which one allele is WT and the other displays a mutation. It can also be due to colony mosaicism where a colony contains a mixture of cells with WT and mutant alleles due to mutations occurring after transformed cells have started to divide. Both mono-allelic mutants and mosaic colonies have been observed in *P. tricornutum* [15, 18].

To determine if the colonies were mosaic or mono-allelic, cells from mutant clones producing mixed PCR products were spread onto selective plates to isolate single sub-clones. Thirty four sub-clones from each clone were screened by PCR (a few examples are presented in Figure 2). Two clones (M2 and M3) were mosaic with a mixture of sub-clones showing either a single band corresponding to the expected deletion (61.5% and 25%, respectively), two bands associated with the WT and expected deletion (25.5% and 28.1%, respectively) or a single band corresponding to the WT urease fragment (13% and 46.9% respectively). For each of the two clones PCR amplicons from three putative bi-allelic sub-clones were sequenced (Figure 2). Four out of six (M2_9, M2_10, M3_10 and M3_11) showed the expected 37nt ‘clean’ deletion without any additional mutations. Precise deletions, such as this, using 2 sgRNAs have previously been generated with high efficiency [37, 43], and allow a large degree of control over the mutation. Two of the sub-clones (M3_9 and M2_12) showed one allele with the expected 37nt deletion and the other with an additional deletion at the 2^nd^ sgRNA cut site. In addition, M2_12 showed a C->G SNP within the sgRNA1 target site. Sub-clones derived from the M1 clone showed WT and 4nt deletion PCR amplicons as seen in the original clone, suggesting that this clone may have a mono-allelic mutation.

Using CRISPR-Cas with one sgRNA can introduce a variety of indels into a locus of interest via the error-prone NHEJ DNA repair mechanism [15]. Cas9 preferentially cuts DNA three nucleotides upstream of the PAM sequence in the seed region [44] and the NHEJ mechanism either repairs a double strand break perfectly or indels are introduced. If cut sites are not cleaved at the same time, when using two sgRNAs, mutations at each site rather than removal of the fragment in between target sites may occur [37]. In this study, however, we report a high occurrence of bi-allelic mutants with precise deletions between the CRISPR-Cas cut sites, suggesting that the Cas9/sgRNA complex is cutting efficiently and DNA ends tend to be repaired perfectly. This allows control over the introduced mutations and gives the chance to avoid introducing in-frame indels.

Restriction digest (results not shown) and sequencing (Figure 2) demonstrated loss off the BccI site in all knock-out clones and HpaII in M2_12 and M1 as a deletion downstream of the cut site is required to remove the HpaII site. This demonstrates that restriction screening can be a valuable tool, however is this case screening for a deletion based band shift by PCR was an efficient way of identifying bi-allelic mutants especially given the limited sgRNA/restriction site interactions available for this gene.

As well as clones from plate selection, one culture from liquid selection (LM1; population of cells transferred to liquid selective media after transformation), showed a single band associated with the bi-allelic 37nt deletion following PCR. This was confirmed by sequencing (Figure 2). PCR screening following growth of LM1 in urea showed only the lower MW band product (results not shown), giving further evidence for a bi-allelic mutation from a population of cells. As small volumes of cells are transferred to fresh media when passaging this may have isolated bi-allelic mutants.

### Growth experiments with mutants

Urease catalyses the breakdown of urea to ammonia allowing it to be used as a source of nitrogen [45]. Sub-clones from different cell-lines with 37 or 38nt deletions were tested for knock-out of the urease gene by looking for a lack of growth when supplemented with urea as the sole nitrogen source.

Cells were nitrogen starved and then transferred to media with either nitrate or urea. Cell counts, cells size and Fv/Fm were measured daily for 7 days. Negative controls to account for any background nitrate in the media were also run in which no nitrate or urea was added for WT cultures.

Four putative bi-allelic mutants (LM1, M4, M2_10 and M3_9) were tested along with WT and the mono-allelic M1_10 over two growth curve experiments. Both LM1 from liquid selection (p=0.0029) and the sub-clone M3_9 (p=0.0000001) showed a significant decrease in growth rate in urea compared to nitrate (Figure 3) as well as a significant 13-18‥ decrease in cell size (Figure 4; p=0.0029 and p=0, respectively). The latter was also apparent with light microscopy (results not shown). Mutants in urea could be easily discerned even without cell counts, as cultures appeared much paler in colour. M4 did not show a difference in growth rate but did show a significant decrease in cell size (p=0.038).The mono-allelic mutant M1_10, displayed higher growth in urea and similar growth to the WT control (Figure 3). This correlates with results from Weyman et al. [17] which showed that despite a reduced protein concentration, a mono-allelic urease knock-out was able to grow in urea. M2_10 which screened as a bi-allelic mutant prior to growth experiments showed a smaller but still significant decrease in growth rate (p=0.0014) (Figure 3) and cell size (p=0.0039) (Figure 4). PCR screening of the urease gene following growth in nitrate and urea showed the expected bi-allelic mutation for LM1, M3_9 and M4, however M2_10 also showed a faint WT band in nitrate and a strong WT band in urea (Figure 5). This suggests that M2_10 was mosaic, with cells containing a functional urease out-competing those with a mutant urease. Given that only a faint WT band was present after growth in nitrate this suggests that the majority of the cells from the sub clone contained the mutant urease, initially accounting for the majority of growth and resulting in a lower but still significant decrease in growth rate.

**Figure 3.**
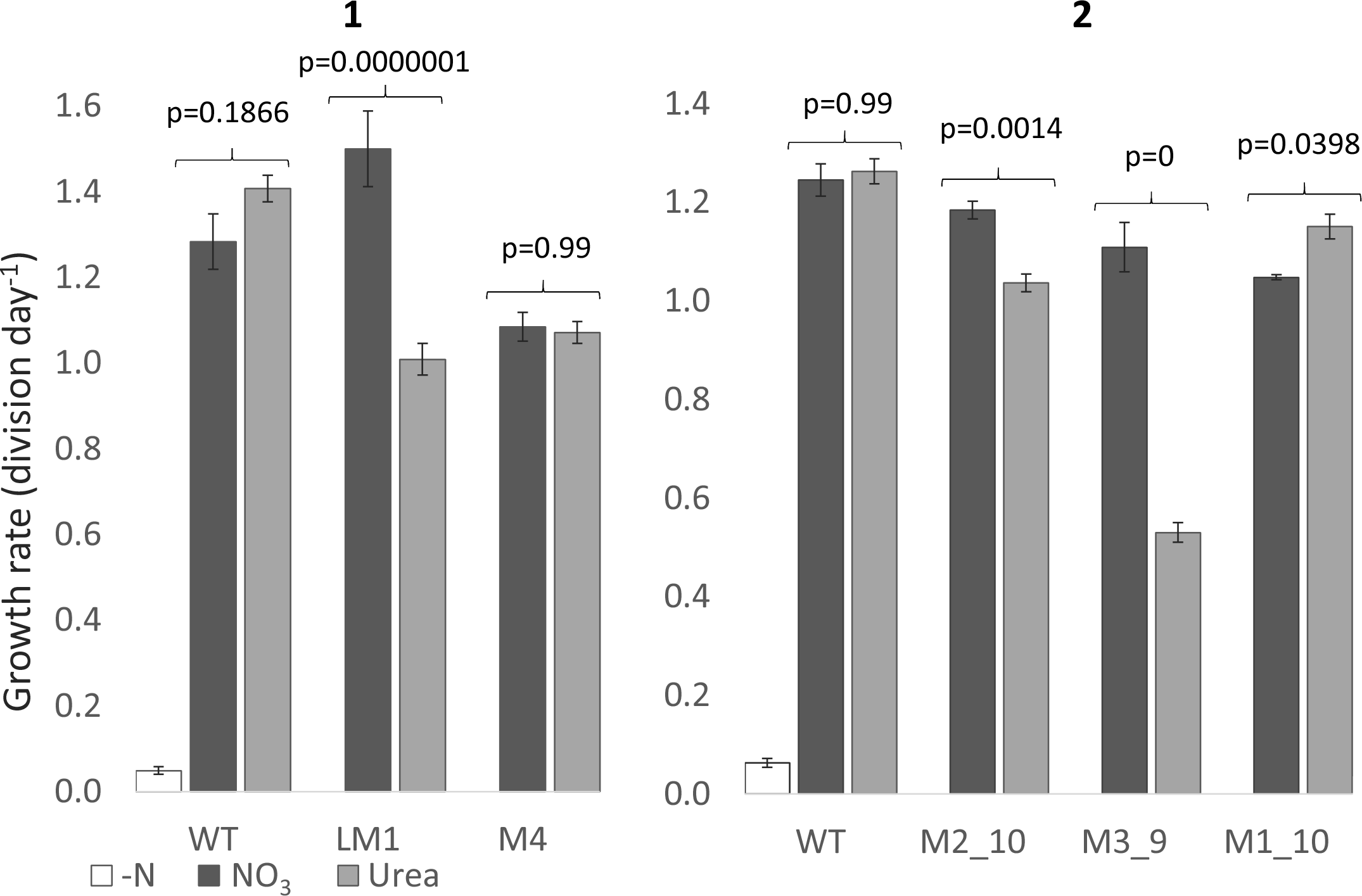
Growth rate of WT and mutant urease cell lines from two separate growth experiments (**1**) & (**2**). The WT cell line was grown in nitrate free (white), nitrate (dark grey) and urea (light grey) enriched media. Mutant cell lines were grown in nitrate or urea enriched media. Growth rate (division day^−1^) was measured in exponential phase and rates compared using analysis of variance with Tukey’s pairwise comparisons.

**Figure 4.**
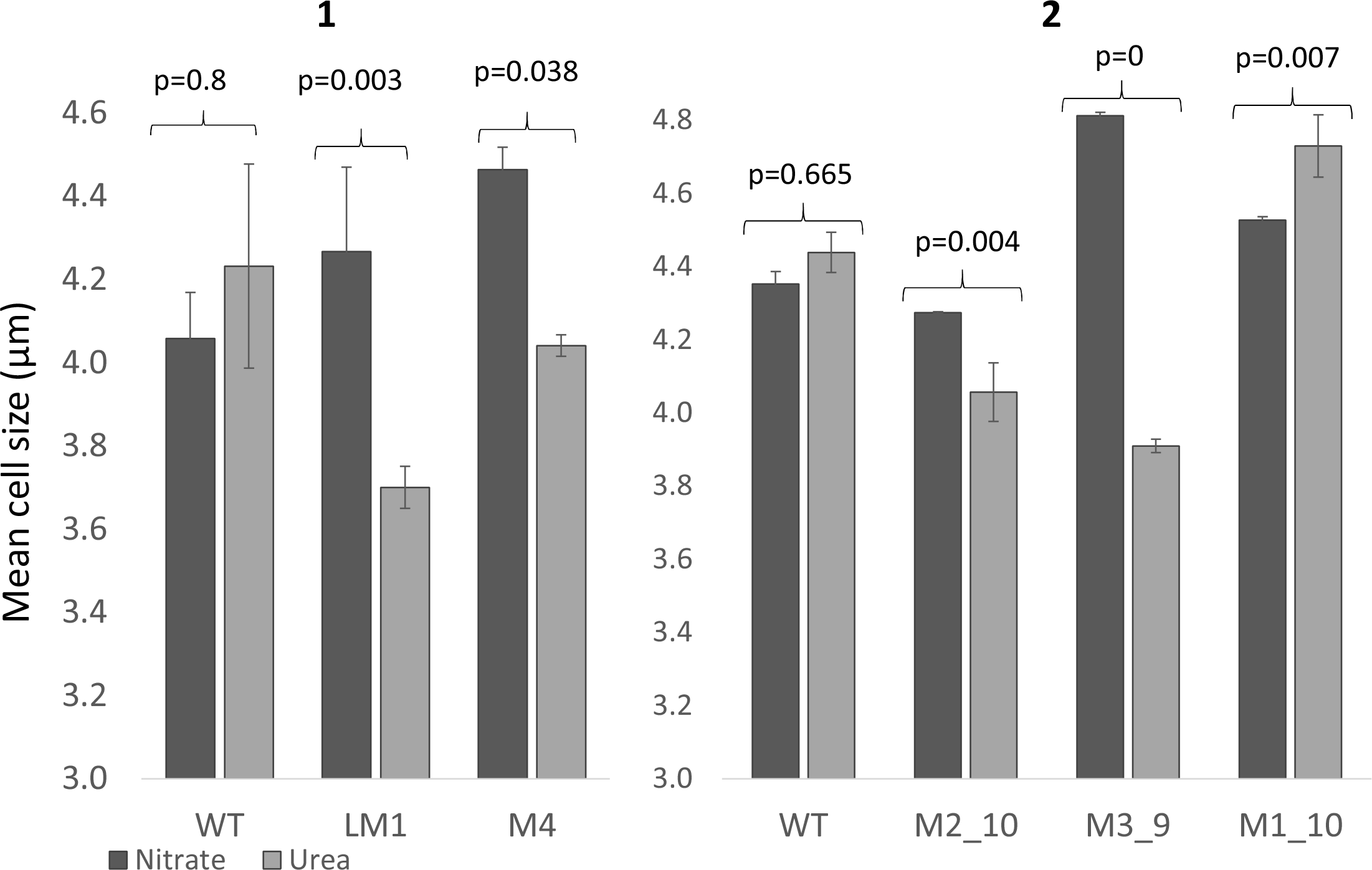
Mean cell size (μm) measured at the end of exponential phase for WT and mutant cultures across two growth experiments (**1**) & (**2**). Cells were grown with nitrate (dark grey) or urea (light grey) as the sole nitrogen source. Cell size was compared using analysis of variance with Tukey’s pairwise comparisons.

**Figure 5.**
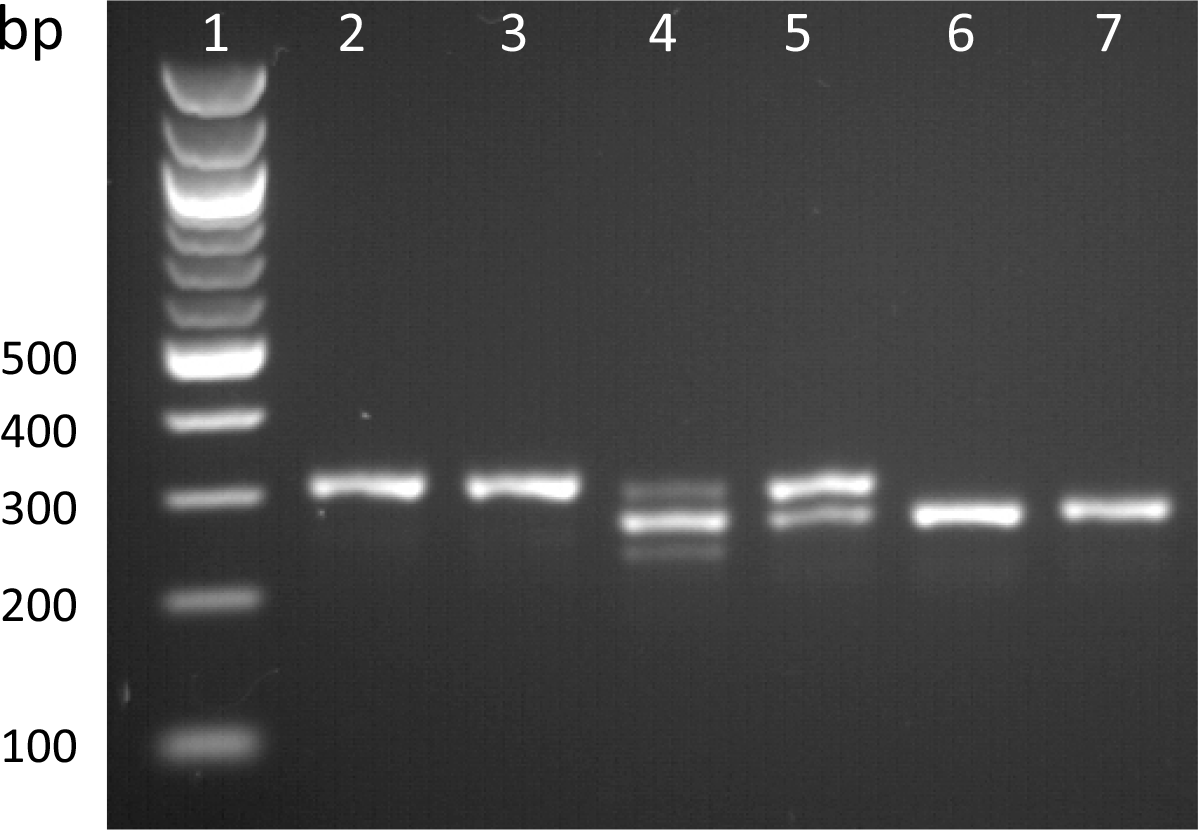
PCR of the targeted urease fragment following growth of WT and mutant cell lines in nitrate or urea. NEB 100bp ladder (**1**), WT in nitrate (**2**) and urea (**3**), M2_ 10 in nitrate (**4**) and urea (**5**) and M3_9 in nitrate (**6**) and urea (**7**).

Knock-out of the urease gene in the diatom *P. tricornutum* prevents growth in urea [17]. Urease mutants in this study still grew in urea but with a lower growth rate and reduced cell-size, characteristics which are associated with nitrogen limitation in diatoms [46, 47] rather than nitrogen starvation. Mutant cell-lines in urea grew to the same density as the same cell-lines in nitrate, but at a lower rate (Figure 3). As nitrogen is an essential nutrient for growth, this suggests that mutant cells in urea still have access to nitrogen, but lower growth rates and cell-size indicates that nitrogen may not be as readily available compared to cells grown with nitrate. Controls in nitrogen free media showed very little growth which suggests that growth of mutants in urea was not due to residual nitrate in the culture. It is unlikely that random integration of the CRlSPR-Cas plasmid is responsible for reduced growth rate in mutants as all four individual mutant cell-lines display increased growth rates when grown in nitrate. Therefore it seems likely that impaired growth of urease mutants in urea is due to a reduction in function of the urease gene.

There are a few possible reasons why a mutation in the urease gene appears to lead to nitrogen limitation rather than nitrogen starvation as seen in *P. tricornutum*. Cells may be able to access nitrogen from another source, separate to the breakdown of urea via urease. Some algae have an alternative pathway for breakdown of urea but this has only been found in Chlorophyceae [48] and blast searches show no evidence of urea carboxylase or allophanate hydroxylase, the enzymes involved in this pathway, in *T. pseudonana*.

The urease gene may still be active but with lower functionality. In *T. pseudonana* urease is modelled to be 807 amino acids. Urease consists of multimers of three sub-units: gamma, beta and alpha, which in TP are translated as one protein. The alpha sub-unit contains the active site which catalyses the breakdown of urea to ammonia [45]. The gamma subunit has no known enzymatic function [49] but may play a role in quaternary structure and stability [45, 50].

Translations of urease sequences with both 37 and 38nt deletions show frame shifts and early stop codons after the deletion in the gamma sub-unit, leading to major disruption of the gamma sub-unit, nonsense down-stream and short products of 24 or 44 amino acid residues (Figure 6). Since all mono-clonal bi-allelic mutants tested for growth in urea had either two alleles with a 37nt deletion or both a 37 and 38nt deletion, it was predicted that the urease gene would no longer be functional. However, several mechanisms exist in eukaryotes which can allow translation of the protein from start codons later in the coding region. These include leaky initiation, re-initiation of ribosomes and internal ribosome entry sites (IRES) [51]. IRES have been shown to become active in yeast following amino acid starvation [51]. If an in-frame translation can occur after the deletion at an IRES or via a mechanism such as re-initiation then the active site located in the alpha-subunit could still be present. The first in-frame ATG after the deletion would start translation of the protein just before the beta sub-unit, leading to an N-terminal truncated protein without the gamma sub-unit but with both the beta and alpha sub-units (Figure 6). Earlier start codons are predicted to result in non-sense and early stop codons.

The 5’ end of the urease coding region was targeted to induce a frame shift and disrupt the protein early on, however it may be better to target the active site or entirely remove the gene. Precise deletions larger than a gene using CRlSPR-Cas and two sgRNAs have been previously demonstrated [43].

**Figure 6.**
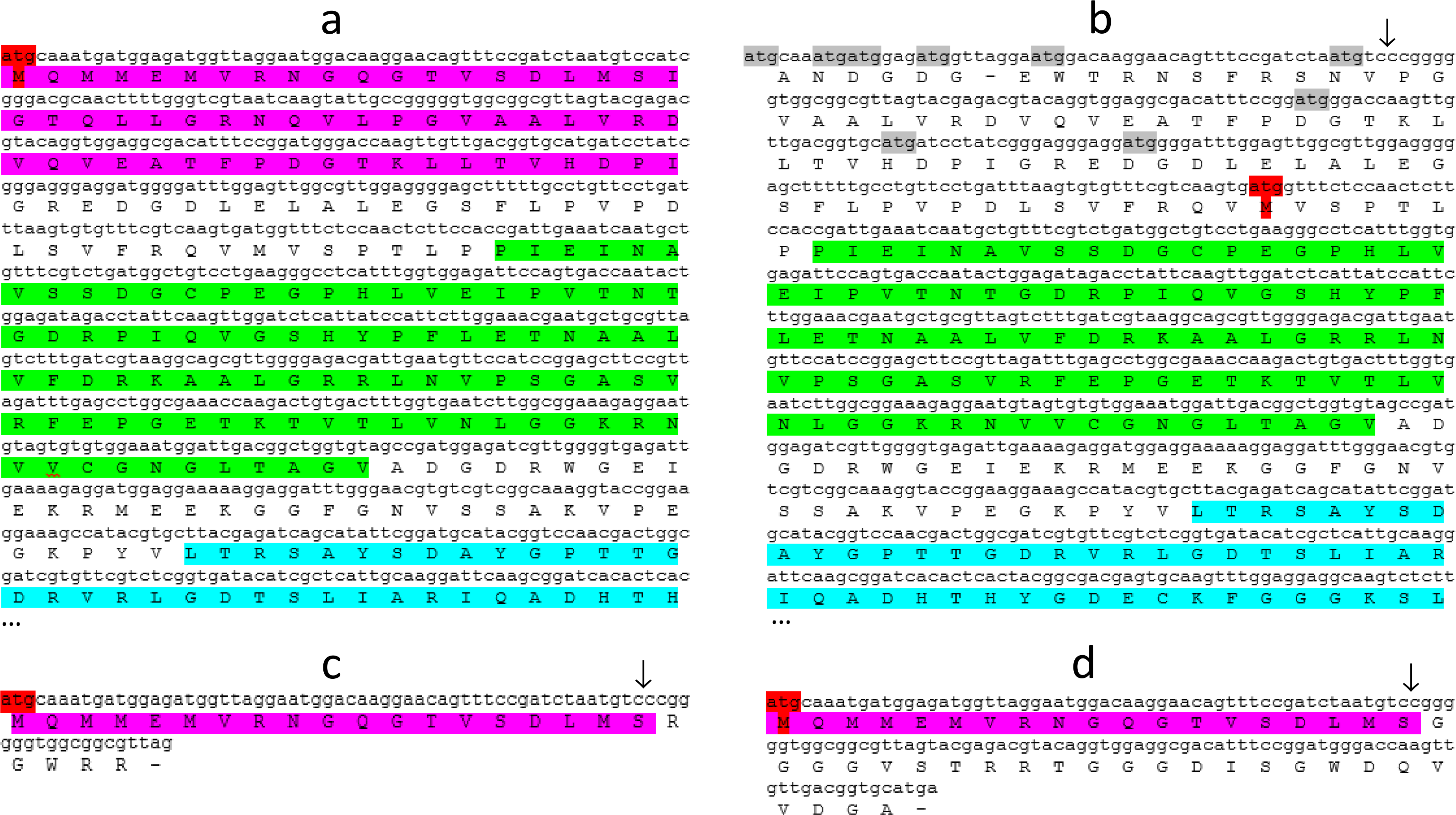
Translated WT urease (**a**), frame 3 (**b**) and frame 1 of urease with the expected 37nt deletion (**c**) and frame 1 of urease with a 38nt deletion (**d**). Position of deletion indicated by ↓. The model WT protein contains 807aa. The figure shows the initial 260 amino acids for **a** and **c** including the start of the alpha sub-unit. Translations are identical for the unshown segments. Gamma (pink), Beta (green) and Alpha (blue) sub-units are highlighted in order. Expected start codon (red) and upstream out-of-frame start codons (grey) are highlighted.

## Conclusions

CRISPR-Cas can precisely and efficiently edit the genome of the diatom *Thalassiosira pseudonana*. Twelve percent of initial colonies and 100% which screened positive for Cas9 showed evidence of a mutation in the urease gene, with many sub-clones showing precise bi-allelic 37nt deletions from two sgRNA DSBs. Screening for the deletion by PCR allowed efficient identification of bi-allelic mutants and Golden Gate cloning allowed easy assembly of a plasmid for CRISPR-Cas. This included adapting the system for *T. pseudonana* by including endogenous promoters and two specific sgRNAs. Due to the flexible modular nature of the cloning system, this can be easily adapted for other genes in *T. pseudonana*. A variety of available online tools were used to design two sgRNAs that would target the early coding region of the urease gene. A reduced growth rate and cell-size phenotype was seen in mutant cell-lines grown in urea compared to nitrate, suggesting that function of the urease may have been impaired rather than removed or an alternative source of nitrogen was available.

As potentially the most important tool in gene editing to date, CRISPR-Cas is fast becoming a key method in the molecular toolbox for a large variety of organisms. This efficient method has huge potential for future work from both an ecological and biotechnology perspective in *T. pseudonana* and can potentially be easily adapted for many other algal species.

## Acknowledgements

This work has been funded by a PhD studentship from the Natural Environment Research Council (NERC) awarded to AH. TM was partially supported by NERC (NE/K013734/1) and the School of Environmental Sciences at UEA. We are thankful to Lewis Dunham for his contributions to the 5’ RACE for identifying the U6 promoter.

---

**Supplementary Figure 1.**
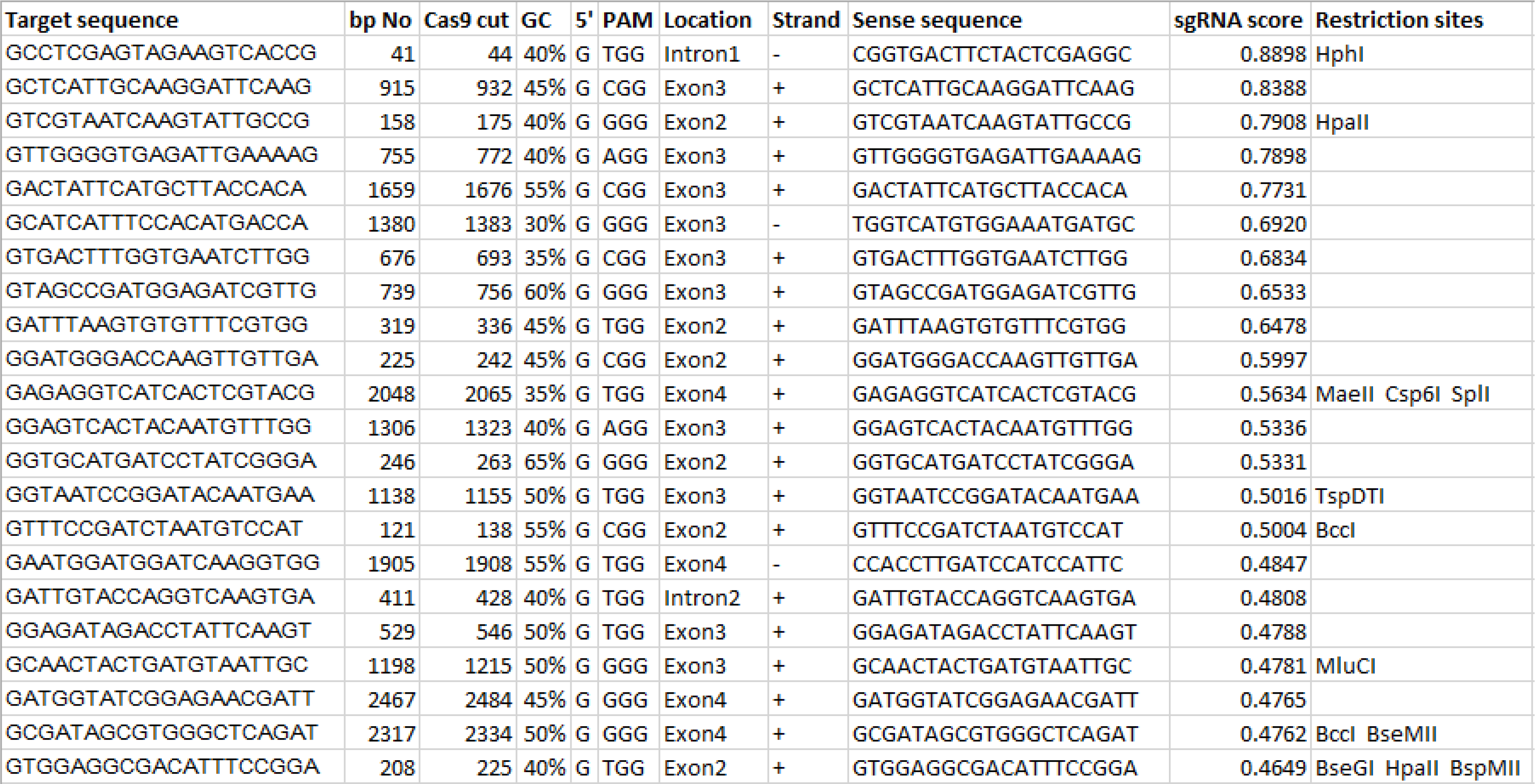
Screenshot of the final spreadsheet for choosing sgRNAs.

## References

1. Kooistra WHCF, Medlin LK: Evolution of the Diatoms (Bacillariophyta). Mol Phylogenet Evol 1996, 6:391–407.

2. Armbrust, E Virginia and Berges, John A and Bowler, Chris and Green, Beverley R and Martinez, Diego and Putnam, Nicholas H and Zhou, Shiguo and Allen, Andrew E and Apt, Kirk E and Bechner M and others: Lawrence Berkeley National Laboratory Lawrence Berkeley National Laboratory. 2004.

3. Bowler C, Allen AE, Badger JH, Grimwood J, Jabbari K, Kuo A, Maheswari U, Martens C, Maumus F, Otillar RP, Rayko E, Salamov A, Vandepoele K, Beszteri B, Gruber A, Heijde M, Katinka M, Mock T, Valentin K, Verret F, Berges J a, Brownlee C, Cadoret J-P, Chiovitti A, Choi CJ, Coesel S, De Martino A, Detter JC, Durkin C, Falciatore A, et al.: The Phaeodactylum genome reveals the evolutionary history of diatom genomes. Nature 2008, 456:239–244.

4. Smetacek V: Diatoms and the ocean carbon cycle. Protist News 1999, 150:25–32.

5. Field CB: Primary Production of the Biosphere: Integrating Terrestrial and Oceanic Components. Science(80-) 1998, 281:237–240.

6. Falkowski PG, Raven JA: Aquatic Photosynthesis. Second. Princeton University Press; 2007.

7. Dolatabadi JEN, de la Guardia M: Applications of diatoms and silica nanotechnology in biosensing, drug and gene delivery, and formation of complex metal nanostructures. TrAC-Trends Anal Chem 2011, 30:1538–1548.

8. Delalat B, Sheppard VC, Rasi Ghaemi S, Rao S, Prestidge C a., McPhee G, Rogers M-L, Donoghue JF, Pillay V, Johns TG, Kröger N, Voelcker NH: Targeted drug delivery using genetically engineered diatom biosilica.Nat Commun 2015, 6:8791.

9. d’Ippolito G, Sardo A, Paris D, Vella FM, Adelfi MG, Botte P, Gallo C, Fontana A: Potential of lipid metabolism in marine diatoms for biofuel production. Biotechnol Biofuels 2015, 8:28.

10. Jeffryes C, Campbell J, Li H, Jiao J, Rorrer G: The potential of diatom nanobiotechnology for applications in solar cells, batteries, and electroluminescent devices. Energy Environ Sci 2011, 4:3930.

11. Kuczynska P, Jemiola-Rzeminska M, Strzalka K: Photosynthetic pigments in diatoms. Mar Drugs 2015, 13:5847–5881.

12. Lander ES: The Heroes of CRISPR. Cell 2016, 164:18–28.

13. Sander JD, Joung JK: CRISPR-Cas systems for editing, regulating and targeting genomes. Nat Biotechnol 2014, 32:347–55.

14. Doudna J a., Charpentier E: The new frontier of genome engineering with CRISPR-Cas9. Science (80-) 2014, 346:1258096–1258096.

15. Nymark M, Sharma AK, Sparstad T, Bones AM, Winge P: A CRISPR/Cas9 system adapted for gene editing in marine algae. Sci Rep 2016, 6(April):24951.

16. Shin S-E, Lim J-M, Koh HG, Kim EK, Kang NK, Jeon S, Kwon S, Shin W-S, Lee B, Hwangbo K, Kim J, Ye SH, Yun J-Y, Seo H, Oh H-M, Kim K-J, Kim J-S, Jeong W-J, Chang YK, Jeong B: CRISPR/Cas9-induced knockout and knock-in mutations in Chlamydomonas reinhardtii. Sci Rep 2016, 6(April):27810.

17. Weyman PD, Beeri K, Lefebvre SC, Rivera J, Mccarthy JK, Heuberger AL, Peers G, Allen AE, Dupont CL: Inactivation of Phaeodactylum tricornutum urease gene using transcription activator-like effector nuclease-based targeted mutagenesis. Plant Biotechnol J 2015, 13:460–470.

18. Daboussi F, Leduc S, Maréchal A, Dubois G, Guyot V, Perez-Michaut C, Amato A, Falciatore A, Juillerat A, Beurdeley M, Voytas DF, Cavarec L, Duchateau P: Genome engineering empowers the diatom Phaeodactylum tricornutum for biotechnology. Nat Commun 2014, 5(May):3831.

19. Poulsen N, Chesley PM, Kröger N: Molecular genetic manipulation of the diatom Thalassiosira pseudonana (Bacillariophyceae). J Phycol 2006, 42:1059–1065.

20. Karas BJ, Diner RE, Lefebvre SC, McQuaid J, Phillips APR, Noddings CM, Brunson JK, Valas RE, Deerinck TJ, Jablanovic J, Gillard JTF, Beeri K, Ellisman MH, Glass JI, Hutchison III C a., Smith HO, Venter JC, Allen AE, Dupont CL, Weyman PD: Designer diatom episomes delivered by bacterial conjugation. Nat Commun 2015, 6:6925.

21. Cook O, Hildebrand M: Enhancing LC-PUFA production in Thalassiosira pseudonana by overexpressing the endogenous fatty acid elongase genes. J Appl Phycol 2015, 28:897–905.

22. Doan TTY, Sivaloganathan B, Obbard JP: Screening of marine microalgae for biodiesel feedstock. Biomass and Bioenergy 2011, 35:2534–2544.

23. Malviya S, Scalco E, Audic S, Vincent F, Veluchamy A, Bittner L, Poulain J, Wincker P, ludicone D, de Vargas C, Zingone A, Bowler C: Insights into global diatom distribution and diversity in the world’ s ocean. Proc Natl Acad Sci 2015, 348:in review.

24. Shrestha RP, Hildebrand M: Evidence for a regulatory role of diatom silicon transporters in cellular silicon responses. Eukaryot Cell 2015, 14:29.

25. Scheffel A, Poulsen N, Shian S, Kröger N: Nanopatterned protein microrings from a diatom that direct silica morphogenesis. Proc Natl Acad Sci U S A 2011, 108:3175–3180.

26. Poulsen N, Scheffel A, Sheppard VC, Chesley PM, Kroger N: Pentalysine clusters mediate silica targeting of silaffins in Thalassiosira pseudonana. J Biol Chem 2013, 288:20100–20109.

27. Xing H-L, Dong L, Wang Z-P, Zhang H-Y, Han C-Y, Liu B, Wang X-C, Chen Q-J: A CRISPR/Cas9 toolkit for multiplex genome editing in plants. BMC Plant Biol 2014, 14:327.

28. Weber E, Engler C, Gruetzner R, Werner S, Marillonnet S: A modular cloning system for standardized assembly of multigene constructs. PLoS One 2011, 6.

29. Doench JG, Hartenian E, Graham DB, Tothova Z, Hegde M, Smith I, Sullender M, Ebert BL, Xavier RJ, Root DE: Rational design of highly active sgRNAs for CRISPR-Cas9-mediated gene inactivation. Nat Biotechnol 2014, 32:1262–7.

30. Xiao A, Cheng Z, Kong L, Zhu Z, Lin S, Gao G, Zhang B: CasOT: A genome-wide Cas9/gRNA off-target searching tool. Bioinformatics 2014, 30:1180–1182.

31. Tarleton R, Peng D: EuPaGDT: a web tool tailored to design CRISPR guide RNAs for eukaryotic pathogens. Microb Genomics 2015, 1:1–7.

32. Nekrasov V, Staskawicz B, Weigel D, Jones JDG, Kamoun S: Targeted mutagenesis in the model plant Nicotiana benthamiana using Cas9 RNA-guided endonuclease. Nat Biotechnol 2013, 31:691–693.

33. Belhaj K, Chaparro-Garcia A, Kamoun S, Nekrasov V: Plant genome editing made easy: targeted mutagenesis in model and crop plants using the CRISPR/Cas system. Plant Methods 2013, 9:39.

34. Price, N.M., Harrison, G.I., Hering, J.G., Hudson, R.J., Nirel, P.M., Palenik, B. and Morel FM: Preparation and Chemistry of the Artificial Algal Culture Medium Aquil. Biol Oceanogr 1989, 6:443–461.

35. Pinto FL, Lindblad P: A guide for in-house design of template-switch-based 5??? rapid amplification of cDNA ends systems. Anal Biochem 2010, 397:227–232.

36. Stothard P: The Sequence Manipulation Suite: JavaScript Programs for Analyzing and Formatting Protein and DNA Sequences. Biotechniques 2000, 28:1102–1104.

37. Brooks, C., Nekrasov, V., Lippman, Z.B. and Van Eck J: Efficient Gene Editing in Tomato in the First Generation Using the Clustered Regularly Interspaced Short Palindromic Repeats/CRISPR-Associated9 Systeml. Plant Physiol 2014, 166:1292–1297.

38. Jacobs TB, LaFayette PR, Schmitz RJ, Parrott W a: Targeted genome modifications in soybean with CRISPR/Cas9. BMC Biotechnol 2015, 15:16.

39. Sakuma T, Nishikawa A, Kume S, Chayama K, Yamamoto T: Multiplex genome engineering in human cells using all-in-one CRISPR/Cas9 vector system. Sci Rep 2014, 4:5400.

40. Port F, Chen H-M, Lee T, Bullock SL: Optimized CRISPR/Cas tools for efficient germline and somatic genome engineering in Drosophila. Proc Natl Acad Sci U S A 2014, 111:E2967–2976.

41. Weber E, Gruetzner R, Werner S, Engler C, Marillonnet S: Assembly of designer tal effectors by golden gate cloning. PLoS One 2011, 6.

42. Krysiak C, Mazus B, Buchowicz J: Relaxation, linearization and fragmentation of supercoiled circular DNA by tungsten microprojectiles. Transgenic Res 1999, 8:303–306.

43. Zheng Q, Cai X, Tan MH, Schaffert S, Arnold CP, Gong X, Chen CZ, Huang S: Precise gene deletion and replacement using the CRISPR/Cas9 system in human cells. Biotechniques 2014, 57:115–124.

44. Jinek M, Chylinski K, Fonfara I, Hauer M, Doudna J a, Charpentier E: A Programmable Dual-RNA-Guided. 2012, 337(August):816–822.

45. Gupta S, Kathait A, Sharma V: Computational Sequence Analysis and Structure Prediction of Jack Bean Urease. 2015, 3:185–191.

46. Olson RJ, Vaulot D, Chisholm SW: Effects of environmental stresses on the cell cycle of 2 marine phytoplankton species. Plant Physiol 1986, 80:918–925.

47. Li W, Gao K, Beardall J: Interactive Effects of Ocean Acidification and Nitrogen-Limitation on the Diatom Phaeodactylum tricornutum. PLoS One 2012, 7.

48. Fan C, Glibert PM, Alexander J, Lomas MW: Characterization of urease activity in three marine phytoplankton species, Aureococcus anophagefferens, Prorocentrum minimum, and Thalassiosira weissflogii. Mar Biol 2003, 142:949–958.

49. Habel JE, Bursey EH, Rho BS, Kim CY, Segelke BW, Rupp B, Park MS, Terwilliger TC, Hung LW: Structure of Rv1848 (UreA), the Mycobacterium tuberculosis urease ?? subunit. Acta Crystallogr Sect F Struct Biol Cryst Commun 2010, 66:781–786.

50. Jabri E, Andrew Karplus P: Structures of the Klebsiella aerogenes urease apoenzyme and two active-site mutants. Biochemistry 1996, 35:10616–10626.

51. Hellen CUT, Sarnow P: Internal ribosome entry sites in eukaryotic mRNA molecules Internal ribosome entry sites in eukaryotic mRNA molecules. 2001:1593–1612.

